# The transmembrane protein Crb2a regulates cardiomyocyte apicobasal polarity and adhesion in zebrafish

**DOI:** 10.1101/398909

**Authors:** Vanesa Jiménez-Amilburu, Didier Y.R. Stainier

## Abstract

Tissue morphogenesis requires changes in cell-cell adhesion as well as in cell shape and polarity. Cardiac trabeculation is a morphogenetic process essential to form a functional ventricular wall. Here we show that zebrafish hearts lacking Crb2a, a component of the Crumbs polarity complex, display compact wall integrity defects and fail to form trabeculae. Crb2a localization is very dynamic, at a time when other cardiomyocyte junctional proteins also relocalize. Before the initiation of cardiomyocyte delamination to form the trabecular layer, Crb2a is expressed in all ventricular cardiomyocytes colocalizing with the junctional protein ZO-1. Subsequently, Crb2a becomes localized all along the apical membrane of compact layer cardiomyocytes and is downregulated by those delaminating. We show that blood flow and Nrg/ErbB2 signaling regulate these Crb2a localization changes. *crb2a* mutants display a multilayered wall with polarized cardiomyocytes, a unique phenotype. Our data further indicate that Crb2a regulates cardiac trabeculation by controlling the localization of tight and adherens junctions in cardiomyocytes. Importantly, transplantation data show that Crb2a controls trabeculation in a CM-autonomous manner. Altogether, our study reveals a critical role for Crb2a during cardiac development.

**Summary statement:** Investigation of the Crumbs polarity protein Crb2a in zebrafish reveals a novel role in cardiac development via regulation of cell-cell adhesion and apicobasal polarity.

## Introduction

Trabeculation is key to cardiac wall maturation (Sedmera et al., 2000; Liu et al., 2010; Samsa et al., 2013), and defects in this process lead to congenital heart malformations (Jenni et al., 1999; Weiford B. C., 2004; Zhang et al., 2013). Prior to trabeculation, cardiomyocytes (CMs) in zebrafish are organized as a single layer. Complex coordination of different signaling cues between endocardium and myocardium including Nrg/ErbB2 signaling (Gassmann et al., 1995; Lee et al., 1995; Meyer and Birchmeier, 1995; Samsa et al., 2013; Samsa et al., 2015; Rasouli and Stainier, 2017) and mechanical forces like blood flow/contractility (Staudt et al., 2014; Li et al., 2016), promote the delamination of a subset of CMs that seed the trabecular layer. Although recent studies have identified several key regulators of different steps in cardiac development including trabeculation, much remains unknown.

During tissue morphogenesis, changes in the shape of individual cells as well as modulation of the actomyosin cytoskeleton and positioning of tight and adherens junctions translate into changes in the whole tissue (Martin et al., 2010; Gorfinkiel, 2013; Heisenberg and Bellaïche, 2013). Cell polarity is a fundamental feature of several cell types and is involved in developmental processes including proliferation, morphogenesis and migration, all these at play during cardiac trabeculation (Martin-Belmonte and Mostow, 2008; Etienne-Manneville, 2013; Grifoni, 2013; Macara and McCaffrey, 2013; Staudt et al., 2014; Uribe et al., 2018). Polarity complexes including PAR3/PAR6/aPKC (Kemphues et al., 1988), Scribble/DLG/LGL (Bilder et al., 2000) and CRUMBS/PALS1/PATJ (Tepass and Knuest, 1990; Tepass, 1996) are known to interact in order to maintain apicobasal identity, cell shape, and tissue integrity. A recent study has shown that *Prkci* mouse mutants show trabeculation defects (Passer et al., 2016), further supporting the involvement of polarity complexes during cardiac trabeculation. CRUMBS (CRB) is a type I transmembrane protein whose extracellular domain is important for the formation of homophilic interactions at the junction between cells (Letizia et al., 2013; Das and Knust, 2018) and whose intracellular domain is key to maintain the identity of the apical domain (Bulgakova and Knust, 2009). Mutations in *Crb* have been found to affect tissue growth, cytoskeletal rearrangement, junction positioning and stability and establishment of apicobasal polarity in flies, zebrafish and mammals (Grawe et al., 1996; Tepass, 1996; Izaddoost et al., 2002; Omori and Malicki, 2006; Chen et al., 2010; Zou et al., 2012; Alves et al., 2013).

Previously, we have reported that CMs in zebrafish and mouse are polarized in the apicobasal axis and that at least in zebrafish CMs undergo apical constriction and depolarization as they delaminate and seed the trabecular layer (Jiménez-Amilburu et al., 2016; Li et al., 2016; del Monte-Nieto et al., 2018). However, how this polarity is regulated in CMs remains unknown. In this study, we identified a new role for Crb2a, a member of the Crb complex, during cardiac trabeculation in zebrafish. Interestingly, Crb2a localization in compact layer CMs changes from junctional to apical at the onset of trabeculation. In *crb2a* mutants, we observed novel and severe defects in CM arrangement and morphology as well as mislocalization of both tight and adherens junctions. Interestingly, mutant CMs transplanted into wild-type (WT) animals were found exclusively in the trabecular layer. These data suggest that the compromised ability of mutant CMs to integrate into the WT compact layer makes them more prone to delaminate. Altogether, these data further emphasize the key role of apicobasal polarity of CMs and its tight regulation in the process of cardiac wall maturation.

## Results

### Crb2a localization changes during cardiac trabeculation in zebrafish

Of the five *crb* family genes in zebrafish, *crb2a* is the highest expressed in the embryonic heart (data not shown). In order to explore the role of Crb2a during cardiac trabeculation, we first performed Crb2a immunostaining in *Tg(myl7:ras-GFP)* embryonic hearts at different developmental stages (Figure 1). At 50 hpf, before the onset of trabeculation, we observed Crb2a highly enriched at the apical junctions in compact layer CMs (**Fig. 1A-A”**). Once trabeculation has been initiated, at 74 hpf, Crb2a has relocalized to the entire apical membrane of compact layer CMs (**Fig. 1B-B”**). The switch of Crb2a localization in CMs from junctional to apical was also observed in 3D maximum intensity projection images of hearts at 51 (**Fig. 1C, 1E and Movie S1**) and 72 (**Fig. 1D, 1E and Movie S2**) hpf. Notably, delaminated CMs in the trabecular layer do not appear to express Crb2a (**Fig. 1B’-B”**). To confirm the specificity of the Crb2a antibody, we performed Crb2a staining in *crb2a*^-^*/-* hearts and found that the immunostaining was lost (**Fig. S1A-B”**). Next, to further investigate the switch from junctional to apical, we performed immunostaining of Crb2a and the TJ marker ZO-1 at 50 and 79 hpf in the *Tg(-0.2myl7:EGFP-podxl)* line, in which the apical side of CMs is labeled by EGFP (Jimenez-Amilburu et al., 2016) (**Fig. S2**). At 50 hpf, Crb2a colocalizes with ZO-1 at the apical junctions between CMs (**Fig. S2A’-A””**), but not with the apical marker Podocalyxin. However, at 79 hpf, Crb2a immunostaining labels the apical side of compact layer CMs colocalizing with Podocalyxin but no longer with the junctional protein ZO-1 (**Fig. S2B’-B”**). Overall, our results show that Crb2a in compact layer CMs transitions from a prominently junctional to an apical localization when trabeculation starts.

**Figure 1.**
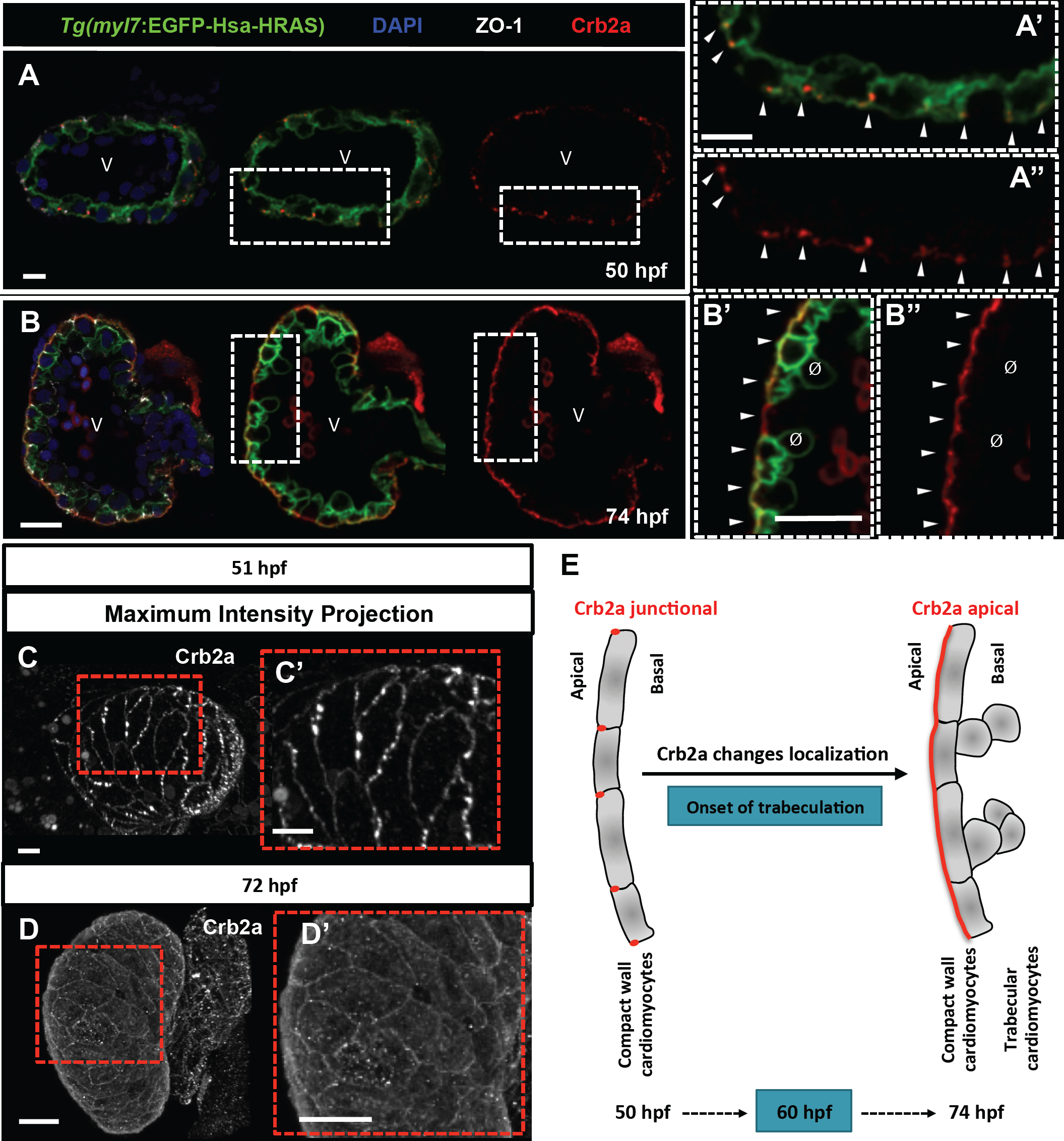
Crb2a localization changes during cardiac trabeculation in zebrafish. (A-B”) Crb2a immunostaining in *Tg(myl7:EGFP-Has-HRAS)* hearts at 50 (A-A”) and 74 (B-B”) hpf. At 50 hpf, Crb2a localizes to the apical junctions between CMs (A’-A”, arrowheads). At 74 hpf, Crb2a immunostaining labels the entire apical membrane of compact layer CMs (B’-B”, arrowheads), and is not observed in delaminated CMs (B’-B”, Ø). (C-D’) Maximum intensity projections of ventricles showing junctional Crb2a staining at 51 (C-C’) and apical Crb2a staining at 72 (D-D’) hpf. (E) Schematic illustration of Crb2a changing its localization in CMs from junctional to apical during the onset of trabeculation. V, ventricle. Scale bars, 20 μm.

### Crb2a localization in developing CMs is modulated by blood flow and Nrg/ErbB2 signaling

In order to investigate what regulates Crb2a localization, we assessed the role of Nrg/ErbB2 signaling and blood flow. As previously reported, blood flow/contractility and Nrg/ErbB2 signaling are essential for cardiac trabeculation (Liu et al., 2010; Peshkovsky et al., 2011; Samsa et al., 2015; Lee et al., 2016; Rasouli and Stainier, 2017). First, to test whether blood flow regulates the localization of Crb2a in CMs, we stopped contractility and blood flow by injecting *tnnt2a* morpholinos (MOs) (Sehnert et al., 2002) into one cell stage embryos. Uninjected larvae at 80 hpf displayed apical localization of Crb2a in compact layer CMs (**Fig. 2A-A”**), while in *tnnt2a* morphants Crb2a was retained at the junctions between compact layer CMs (**Fig. 2B-2B”**). Next, we treated *Tg(myl7:Hras-GFP)* embryos at 54 hpf, i.e., before trabeculation starts, with 10 μM PD168393, an ErbB2 inhibitor, or DMSO as a control. At 96 hpf, in DMSO-treated larvae, Crb2a immunostaining was found along the entire apical membrane of compact layer CMs (**Fig. 2C-2C”, 2E**). Interestingly, larvae treated with the ErbB2 inhibitor at the same stage displayed a significant increase in junctional Crb2a (average of 14.69 compared to 4.57 in DMSO-treated larvae) (**Fig. 2D-D”, E**). Altogether, these data suggest that blood flow and ErbB2 signaling are involved in regulating Crb2a localization in CMs.

**Figure 2.**
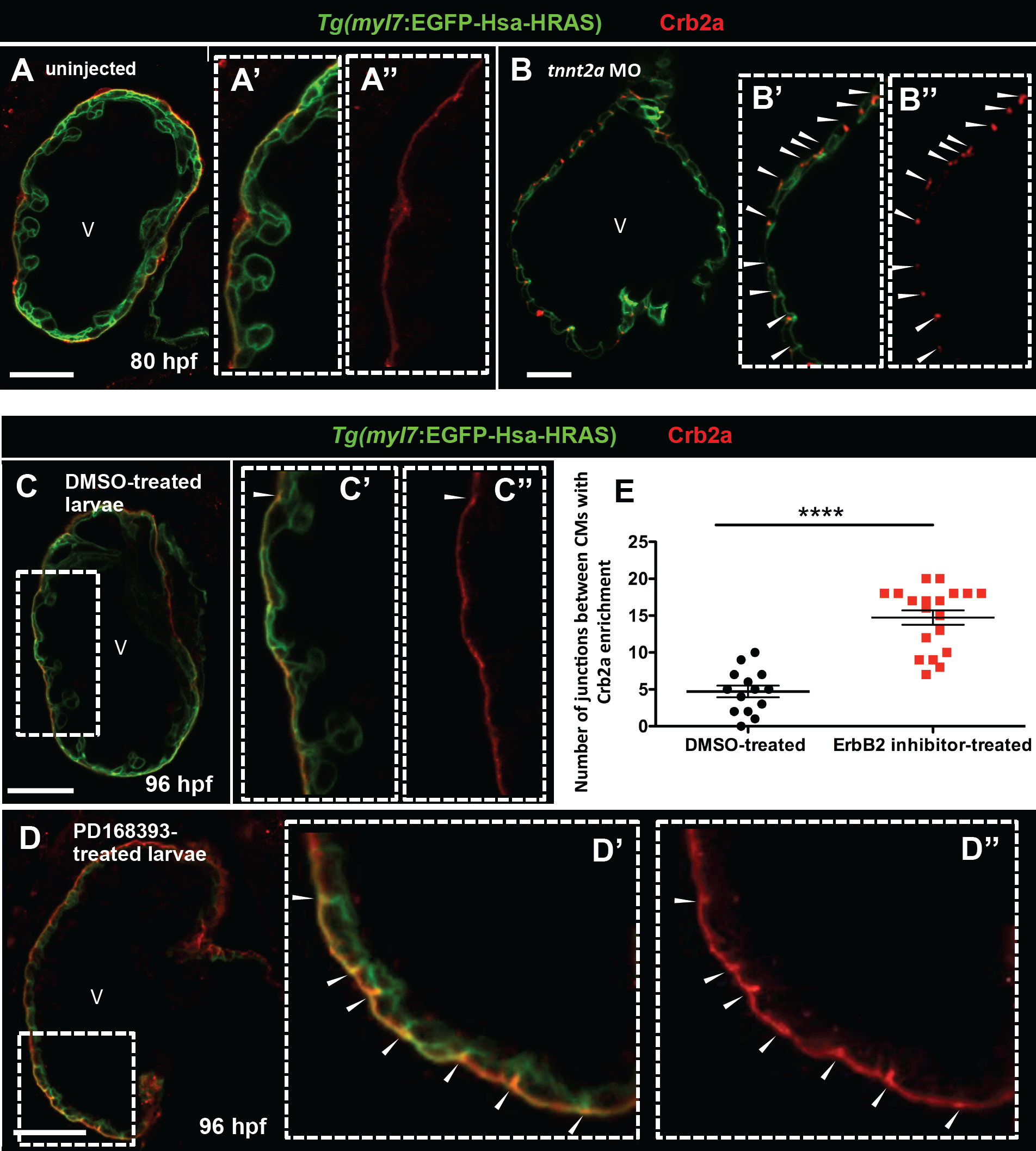
Crb2a localization in developing CMs is modulated by blood flow and Nrg/ErbB2 signaling. (A-B”) Crb2a immunostaining in *Tg(myl7:EGFP-Hsa-HRAS)* control (A) and *tnnt2a* morphant (B) hearts at 80 hpf. Higher magnification images show apical localization of Crb2a in control hearts (A’-A”). Arrowheads in higher magnification images point to junctional localization of Crb2a between CMs in *tnnt2a* morphants (B’-B”). (C-D”) Crb2a immunostaining in 96 hpf *Tg(myl7:EGFP-Hsa-HRAS)* hearts treated between 54 and 96 hpf with DMSO (C-C”) or PD168393 (D-D”). Arrowheads in higher magnification images point to Crb2a enrichment at the junctions between CMs in DMSO-(C’-C”) and PD168393-(D’-D”) treated larvae. (E) Number of junctions between CMs with Crb2a enrichment (DMSO-treated larvae, n=14; ErbB2 inhibitor-treated larvae, n=19). Each dot represents one heart. Data are shown as mean ± SEM. ^****^*P* < 0.0001 by Student’s t test. V, ventricle. Scale bars, 20 μm.

### *crb2a*^-^*/-* hearts display disrupted compact wall integrity and fail to form trabeculae

To investigate the role of Crb2a in the heart during cardiac trabeculation, we studied the *ome*/*crb2a* mutant (Malicki et al., 1996; Omori and Malicki, 2006). First, to characterize CM morphology during cardiac trabeculation, we crossed the *crb2a*^-^*/-* with the CM membrane reporter line *Tg(myl7:MKATE-CAAX).* At 80 hpf, after the onset of trabeculation, *crb2a*^-^*/-* exhibit a more rounded and smaller heart compared to *crb2a*^*+/+*^ (**Fig. 3A-B**). At the same stage, we observed that while *crb2a*^*+/+*^ hearts display a single layer of CMs in the compact wall and few delaminated CMs (**Fig. 3A’-A”, E, F**), the mutant cardiac wall is composed of two or three layers of CMs and appears disorganized (**Fig. 3B’-B”’, E’, F**). In addition, the CMs in the mutant heart seem to be more elongated than those in *crb2a*^*+/+*^ hearts (**Fig. 3A’-A”, B’-B”’**). We imaged the same mutant larvae at 98 hpf, and interestingly observed that they did not exhibit myocardial protrusions into the lumen of the heart and failed to form trabeculae (**Fig. 3C-C’**) while *crb2a*^*+/+*^ larvae at this stage presented a complex trabecular network (**Fig. 3D-D’**). Next, we crossed *crb2a*^-^*/-* with the *Tg(kdrl:EGFP); Tg(myl7:MKATE-CAAX)* lines to analyze whether problems in cardiac jelly degradation might account for this phenotype. We did not observe differences in cardiac jelly thickness between *crb2a*^*+/+*^ (**Fig. S3A-C’**) and *crb2a*^-^*/-* (**Fig. S3B-D’**) animals at any of the developmental stages analyzed. To check whether the multilayering phenotype was present before the onset of trabeculation, we imaged 52 hpf embryos, and found that while *crb2a*^*+/+*^ hearts consist of a single layer of CMs (**Fig. S4A-A’, D**), 86% of the mutant hearts display multilayering (**Fig. S4B-B’, D**) and the other 14% exhibit a single layer of CMs (**Fig. S4C-C’, D**). In order to further study the early cardiac phenotype of these mutants, we imaged them at 36 hpf and found that while *crb2a*^*+/+*^ and *crb2a*^*+/*^- hearts have looped (**Fig. S5A, B**), *crb2a*^-^*/-* hearts display no or delayed looping (**Fig. S5C-C’**). We also analyzed the circulation in these mutants at 48 hpf and found that 40% of the mutants exhibit no or a low number of circulating blood cells, while the other 60% exhibit a WT like circulation (**Fig. S6D**). To evaluate cardiac performance in *crb2a*^-^*/-* hearts, we measured atrial fractional shortening (FS) (**Fig. S6A-C’**). Brightfield images of the beating heart were acquired at 48 hpf and FS was calculated in *crb2a*^*+/+*^and *crb2a*^*+/*^-, as well as in *crb2a*^-^*/-* with and without circulation (Movies S3, S4, S5 and S6). This analysis showed that *crb2a*^-^*/-* without circulation have a significant decrease in FS compared to *crb2a*^*+/+*^ or *crb2a*^*+/*^-, while mutants with circulation present a slight but not significant reduction in FS (**Fig. S6E**). In order to test whether this reduction in contractility was due to sarcomere defects in CMs, we immunostained 80 hpf *Tg(myl7:MKATE-CAAX)*; *crb2a*^-^*/-* hearts for α-actinin, and observed no differences in sarcomeric structure between *crb2a*^*+/+*^ and *crb2a*^-^*/-* (**Fig. S6F-G’**). To avoid any confounding effects, subsequent experiments in this manuscript were performed exclusively with *crb2a-/-* exhibiting WT like circulation. Taken together, these data show that Crb2a plays an important role during cardiac trabeculation.

**Figure 3.**
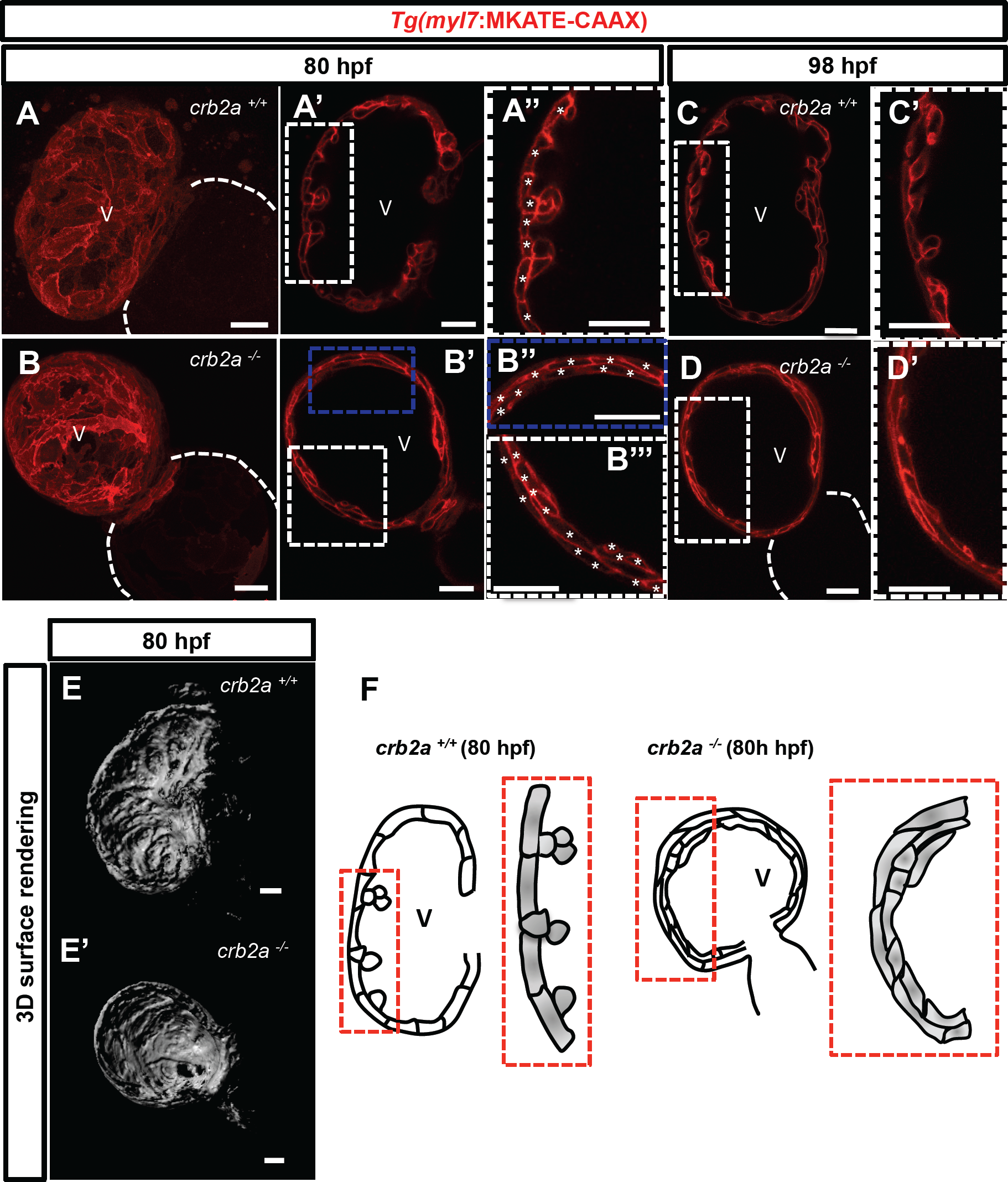
*crb2a*^-^*/-* hearts display disrupted compact wall integrity and fail to form trabeculae. (A-B) Confocal images (maximum intensity projections) of 80 hpf *Tg(myl7:MKATE-CAAX) crb2a*^*+/+*^ (A) and *crb2a*^-^*/-* (B) hearts. (A’-D’) Confocal images (mid-sagittal sections) of *Tg(myl7:MKATE-CAAX)* hearts of *crb2a*^*+/+*^ (A-A” and C-C’) and *crb2a*^-^*/-* (B-B” and D-D’) at 80 and 98 hpf. Asterisks in A” and B” mark individual CMs in compact wall. (E-E’) 3D surface rendering images of 80 hpf *crb2a*^*+/+*^ (E) and *crb2a*^-^*/-* (E’) ventricles. (F) Schematic illustration of myocardial wall in 80 hpf *crb2a*^*+/+*^ and *crb2a*^-^*/-* ventricles. V, ventricle. Scale bars, 20 μm.

### Inner layer of CMs in *crb2a*^-^*/-* are polarized in the apicobasal axis

To gain further insight into the polarization of CMs in *crb2a*^-^*/-* hearts, we crossed these mutants with the *Tg(-0.2myl7:EGFP-podxl)* line. As previously reported, at 80 hpf in *crb2a*^*+/+*^ ventricles, EGFP-Podocalyxin localizes to the apical side of compact layer CMs and covers the entire cortex in delaminated CMs (**Fig. 4A-A’, C**) (Jiménez-Amilburu et al., 2016). In *crb2a*^-^*/-* ventricles, CMs forming the inner layer of the wall were also polarized as per their apical localization of EGFP-Podocalyxin (**Fig. 4B-B’, C**). In an attempt to test whether this multilayering phenotype was due to an increase in CM number, we assessed CM proliferation in mutant hearts. To quantify proliferating CMs we crossed *crb2a*^-^*/-* with *Tg(myl7:nlsdsRED);Tg(myl7:VenusGeminin)* and examined maximum intensity projection images at 80 hpf (**Fig. S7A-C**). We found that the number of total CMs and proliferating CMs was similar in *crb2a*^*+/+*^, *crb2a*^*+/*^-, and *crb2a*^-^*/-* (**Fig. S7D, E**). Taken together, these data show that Crb2a plays a role in modulating CM polarity and specifically to prevent CM delamination from the compact layer in the absence of Nrg/ErbB2 signaling.

**Figure 4.**
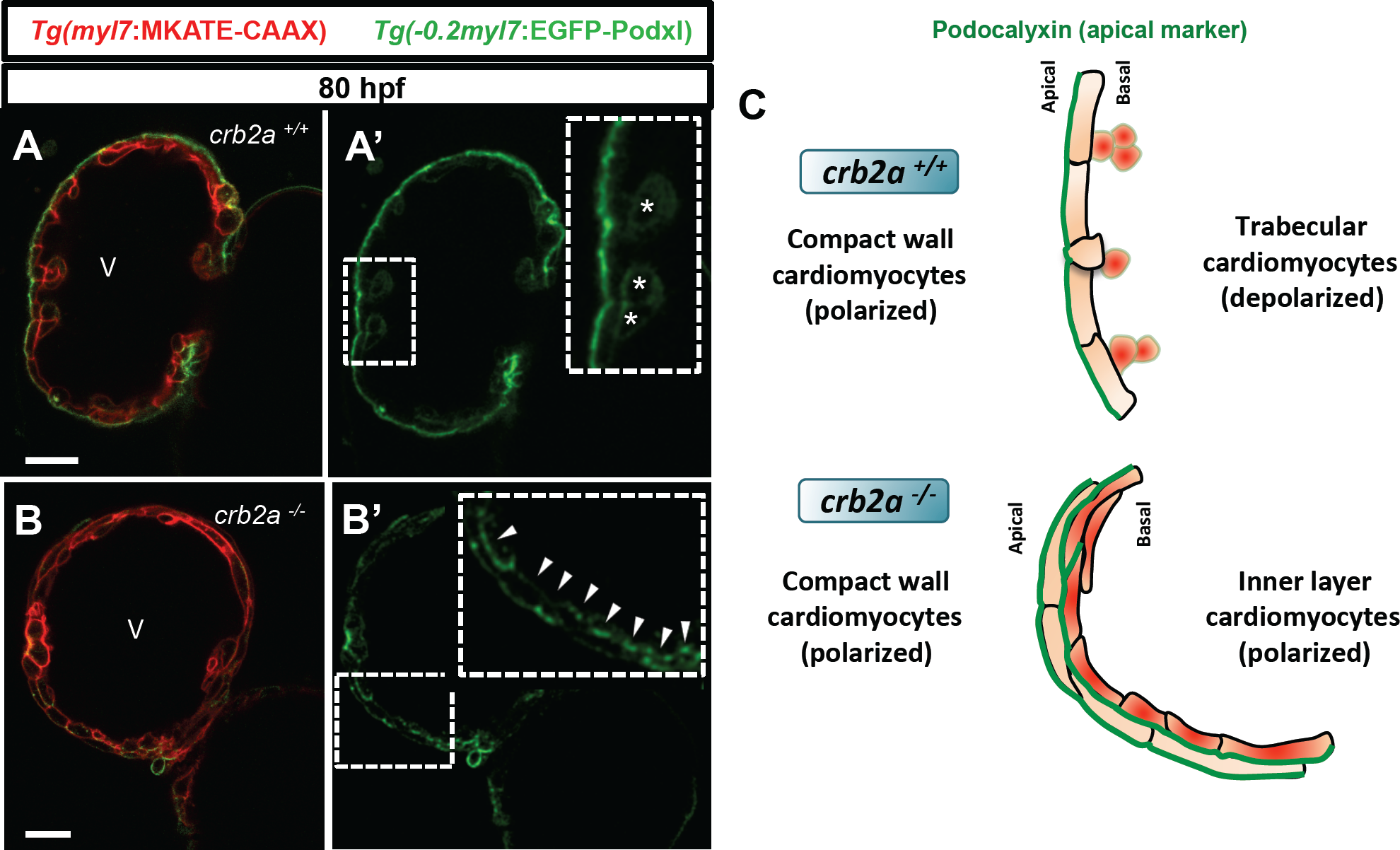
Inner layer of CMs in *crb2a*^-^*/-* are polarized in the apicobasal axis. (A-B’) Confocal images (mid-sagittal sections) of 80 hpf *Tg(-0.2myl7:EGFP-podxl);Tg(myl7:MKATE-CAAX)* hearts of *crb2a*^*+/+*^ (A-A’) and *crb2a*^-^*/-* (B-B’). Asterisks in A’ indicate delaminated CMs and arrowheads in B’ point to polarized CMs. (C) Schematic illustration of A’ and B’ showing depolarized CMs in *crb2a*^*+/+*^ and polarized CMs in the inner layer of *crb2a*^-^*/-*. V, ventricle; At, atrium Scale bars, 20 μm.

### Tight and adherens junctions are mislocalized in *crb2a*^-^*/-* hearts

The Crb complex regulates the localization of tight and adherens junctions during cellular rearrangements (Grawe et al., 1996; Izaddoost et al., 2002). To investigate whether Crb2a regulates tight and adherens junctions during cardiac development, we immunostained embryos of the AJ reporter line *TgBAC(cdh2:cdh2-EGFP)* for Crb2a and ZO-1. At 54 hpf, before the onset of trabeculation, Crb2a is found at the junctions between CMs and colocalizes with ZO-1 (Fig S9A-A””, C). At this stage, N-Cadherin localizes laterally in CMs showing no overlap with Crb2a or ZO-1 (**Fig. S8A-A””, C**). In order to carefully study in vivo TJ dynamics specifically in CMs during cardiac trabeculation, we generated a *Tg(-0.2myl7:ZO-1-EGFP)* line. Using this reporter line we were able to reproduce the endogenous localization of ZO-1 during trabeculation (**Fig. S9**). Once trabeculation initiates and Crb2a becomes apical in compact layer CMs, ZO-1 remains junctional and N-cadherin extends baso-laterally in compact layer CMs (**Fig. S8B-B””, C**). Next, we analyzed whether TJ and AJ were affected in *crb2a*^-^*/-* hearts. First, we performed immunostaining for ZO-1 in *Tg(myl7:MKATE-CAAX); crb2a*^-^*/-* at 79 hpf. Maximum intensity projections and single plane images of *crb2a*^*+/+*^ larvae show junctional localization of ZO-1 between compact layer CMs (**Fig. 5A-A’, B-B’**). However, *crb2a*^-^*/-* hearts at the same stage display mislocalization of ZO-1 in both compact and inner layer CMs (**Fig. 5A”-A””, B”-B””**). Additionally, we also performed immunostaining for N-cadherin in *Tg(myl7:MKATE-CAAX)*; *crb2a*^-^*/-* larvae at 79 hpf. Analysis of maximum intensity projection images of *crb2a*^-^*/-* ventricles showed that the lateral localization of N-Cadherin between compact layer CMs was lost in mutant hearts (**Fig. 5C-C’, D-D’**), appearing punctate in both apical and basal membranes (**Fig. 5C”-C””, D”, D””**). These data show that junctional reorganization in CMs is likely to play a role during CM delamination and suggest a role for Crb2a in regulating both tight an adherens junctions in CMs during cardiac development.

**Figure 5.**
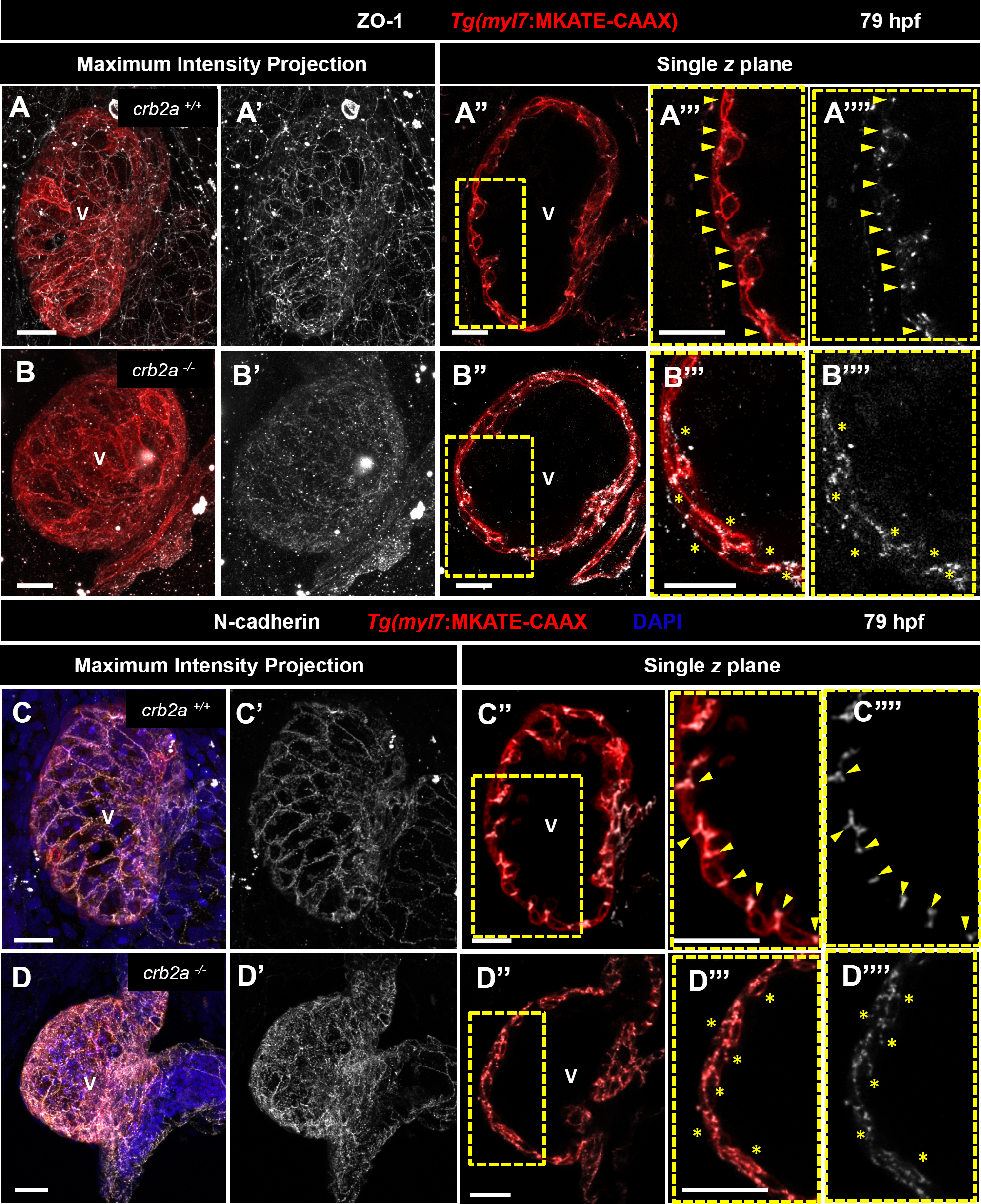
Tight and adherens junctions are mislocalized in *crb2a*^-^*/-* hearts. (A-B””) ZO-1 immunostaining in 79 hpf *Tg(myl7:MKATE-CAAX)* hearts. (A-B’) Maximum intensity projection images show that ZO-1 localizes to the junctions between CMs in *crb2a*^*+/+*^ (A’), but its junctional pattern is fragmented in *crb2a*^-^*/-* (B’). (A”-B”) Confocal images (mid-sagittal sections) of *Tg(myl7:MKATE-CAAX)* hearts. (A”’-B””) High magnifications of single plane images show the junctional localization of ZO-1 between CMs in *crb2a*^*+/+*^ (A”’-A””, arrowheads), while in *crb2a*^-^*/-* ZO-1 appears to outline compact layer CMs at their apical and basal sides (B”’-B””, asterisks). (C-D””) N-Cadherin immunostaining and DAPI in 79 hpf *Tg(myl7:MKATE-CAAX)* hearts. (C-D’) Maximum intensity projection images show that N-Cadherin localizes at the junctions between CMs in *crb2a*^*+/+*^ embryos (C’) but is mislocalized at the junctions of CMs in *crb2a*^-^*/-* (D’). (C”-D”) Confocal images (mid-sagittal sections) of *Tg(myl7:MKATE-CAAX)* hearts. (C”’-D””) High magnifications of single plane images show the lateral localization of N-Cadherin between CMs in *crb2a*^*+/+*^ (C”’-C””, arrowheads), while in *crb2a*^-^*/-* N-Cadherin localization appears more punctate in both the apical and basal membranes of compact layer CMs (D”’-D””, asterisks). V, ventricle. Scale bars, 20 μm.

### Crb2a functions cell-autonomously in CMs during cardiac trabeculation

To gain further insight into Crb2a function during cardiac trabeculation, we generated chimeric embryos by transplanting at the mid-blastula stage *Tg(myl7:MKATE-CAAX) crb2a*^*+/+*^, *crb2a*^*+/*^- and *crb2a*^-^*/-* cells into *Tg(myl7:Hras-GFP) crb2a*^*+/+*^ animals. After transplantation, larvae were imaged at 100 hpf and subsequently genotyped. We found that *crb2a*^*+/+*^ and *crb2a*^*+/*^- CMs in *crb2a*^*+/+*^ ventricles localized in both the compact and trabecular layers. Strikingly, *crb2a*^-^*/-* CMs in *crb2a*^*+/+*^ ventricles were exclusively found in the trabecular layer (**Figures 6A-C**). To extend these observations, we quantified the number of donor-derived CMs in the compact and trabecular layers in chimeric larvae at 100 hpf, and found that 100% of the *crb2a*^-^*/-* CMs populated the trabecular layer of *crb2a*^*+/+*^ ventricles (**Figure 6D**). These data show that in the absence of Crb2a function, CMs leave the compact layer, indicating that Crb2a functions cell-autonomously in CMs.

**Figure 6.**
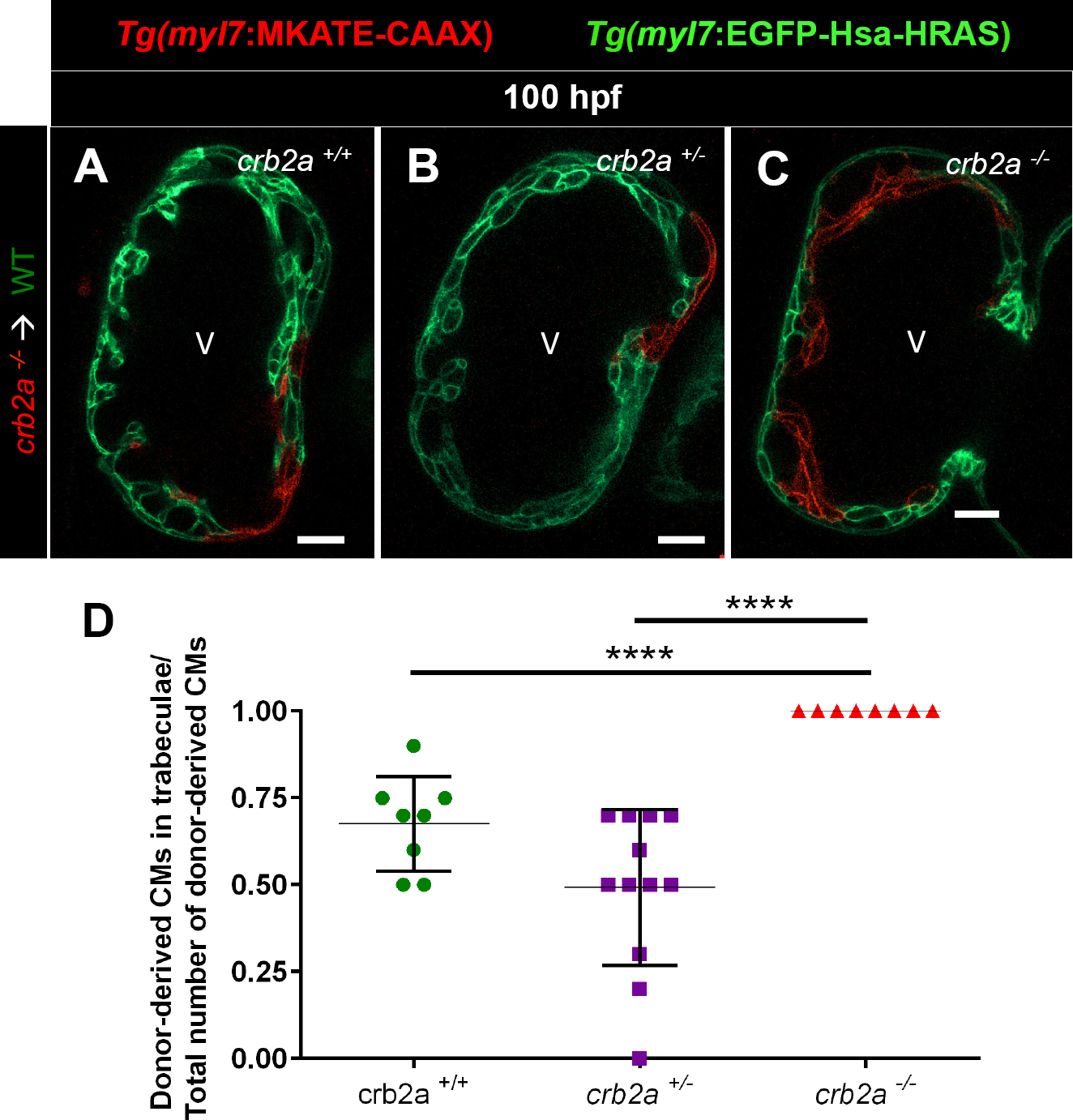
Crb2a functions cell-autonomously in CMs during cardiac trabeculation. (A-C) Confocal images (mid-sagittal sections) of mosaic 100 hpf *crb2a*^*+/+*^ *Tg(myl7:EGFP-Has-HRAS)* host hearts transplanted with *Tg(myl7:MKATE-CAAX) crb2a*^*+/+*^, *crb2a*^*+/*^- or *crb2a*^-^*/-* cells. (D) Percentage of donor-derived CMs in trabeculae versus total number of donor-derived CMs. Each dot represents one heart. Data are shown as mean ± SEM. \*\*\*\**P* < 0.0001 by Student’s t test. V, ventricle. Scale bars, 20 μm.

### Discussion

Cardiac trabeculation in zebrafish has been proposed as an EMT-like process whereby a subset of CMs undergo apical constriction and lose apicobasal polarity as they delaminate to seed the trabecular layer (Liu et al., 2010; Staudt et al., 2014; Cherian et al., 2016; Jiménez-Amilburu et al., 2016). However, it remains unclear how CM apicobasal polarity is regulated. Therefore, we set out to investigate whether and how Crb2a, a member of the Crb polarity complex, regulates cardiac trabeculation. In this study, we present *in vivo* evidence that Crb2a has a key role in maintaining CM apicobasal polarity as well as in regulating junctional rearrangements during early cardiac development.

The coordinated action of different polarity proteins to modulate apicobasal identity, together with cytoskeletal and junctional rearrangements, maintain cell shape and tissue integrity (Hurd et al., 2003; Kempkens et al., 2006). Particularly, Crb has a conserved role in many epithelial tissues, regulating apicobasal polarity and maintaining junctional integrity during morphogenetic processes (Grawe et al., 1996; Izaddoost et al., 2002). Here, we found that the Crb polarity protein Crb2a shifts its localization in CMs from junctional to apical just prior to the onset of trabeculation. Accordingly, CMs have been reported to have a great cellular plasticity during cardiac trabeculation, losing their cobblestone-like shape once they start delaminating (Liu et al., 2010; Staudt et al., 2014). Our observations are consistent with previous reports in *Drosophila* where during follicular morphogenesis, *Crb* is lost from the marginal zone (the most apical surface), relocalizes to the adherens junctions and later reappears at the marginal zone during transitional changes in cell shape (Sherrard and Fehon, 2015; Wu et al., 2016). We found that upon blocking Nrg/ErbB2 signaling, Crb2a in CMs fails to relocalize from the junctions to the apical surface. Similarly, recent data in *Drosophila* show that after depletion of cytoskeletal proteins important for *Crb* localization, apical *Crb* becomes junctional (Sherrard and Fehon, 2015). These data show that Crb2a localization is very dynamic and that Nrg/ErbB2 signaling, which is essential for cardiac trabeculation, is required for its junctional to apical shift.

Zebrafish lacking Crb2a display a compact ventricular wall composed of multiple disorganized layers of CMs. This phenotype is similar to that in *Drosophila* oogenesis where early loss of Crb leads to multilayering (Tanentzapf et al., 2000; Pénalva and Mirouse, 2012). While delaminating CMs in WT hearts lose their apicobasal identity (Jiménez-Amilburu et al., 2016), all CMs in *crb2a* mutants retain it, including those in the inner layers, suggesting that the inner layer CMs in *crb2a* mutants are not trabecular CMs. Furthermore, we observed that *crb2a* mutants display mislocalization of both tight and adherens junctions between CMs. Similar to these data, loss of Crb in epithelial cells in *Drosophila* causes defects in cell-cell adhesion and tissue integrity (Tepass and Knust, 1990; Grawe et al., 1996; van de Pavert et al., 2004; Letizia et al., 2013; Flores-Benitez and Knust, 2015). During epithelial morphogenesis, changes in cell shape and junctional localization require a tight regulation of the connection between junctional proteins and the cytoskeleton (Costa et al., 1998; Raich et al., 1999; Martin et al., 2010). Based on these and other data, we hypothesize that lack of Crb2a causes loss in cell-cell adhesion of CMs.

We observed that *crb2a* mutant CMs in WT ventricles all end up in the trabecular layer. Similarly, *wwtr1* mutant CMs in WT ventricles have a preference to enter the trabecular layer (Lai et al., 2018). In addition, cardiac phenotype of these mutants is similar in the sense that they both exhibit CM multilayering, although it is more pronounced and starts at an earlier stage in *crb2a* mutants. Interestingly, we found that Crb2a immunostaining was mostly gone in *wwtr1* mutant hearts (data not shown), suggesting that this reduction in Crb2a expression is causing the mislocalization of adherens junctions in *wwtr1* mutant hearts (Lai et al., 2018). In summary, the lack of Crb2a functions in zebrafish CMs, affects their ability to remain in a monolayer, subsequently leading to a lack of trabeculae.

## Material and Methods

### Ethics statement

All experiments using zebrafish were performed in accordance with German animal protection laws and approved by the local governmental animal protection committee. Zebrafish were maintained according to standard protocols (http://zfin.org).

### Transgenic and mutant zebrafish lines

Transgenic lines used in this study are as follows: *Tg(-0.2myl7:EGFP-podocalyxin)*^*bns*103^ (Jiménez-Amilburu et al., 2016), abbreviated *Tg(-0.2myl7:EGFP-podxl)*; *Tg(myl7-MKATE-CAAX)*^*sd*11^ (Lin et al., 2012), abbreviated *Tg(myl7:MKATE-CAAX)*; *Tg(myl7:EGFP-Has.HRAS)*^*s*883^ (D’Amico et al., 2007), abbreviated *Tg(myl7:ras-EGFP); Tg(myl7:mVenus-gmnn)*^*ncv^43^Tg*^ (Jiménez-Amilburu et al., 2016), abbreviated *Tg(myl7:mVenus-gmnn); Tg(kdrl:EGFP) s843* (Jin et al., 2005), abbreviated *Tg(kdrl:EGFP); TgBAC(cdh2:cdh2-EGFP,crybb1:ECFP)*^*zf517*^ (Revenu et al., 2014), abbreviated *TgBAC(cdh2:cdh2-ECFP); Tg(-5.1myl7:DsRed2-NLS)*^*f^2^T*^*g* (Rottbauer et al., 2002), abbreviated *Tg(myl7:nlsDsRed).*

Mutant line used in this study is as follows: *crb2a*^*m^598^/m*598^ (Omori and Malicki, 2006) abbreviated *crb2a*^-^*/-*. To screen for adult carriers or discriminate between WT, *crb2a*^*+/*^- and *crb2a*^-^*/-* at embryonic/larval stages we used genomic DNA extracted from clipped fins. PCR was performed using the following primers, PCR: 5’-TCAGGCTTGTCCTTCAAGTC-3’ and 3’-TTACTTGGCTCAGGTGTGTC-5’. The product from PCR1 was then used to perform PCR2 using the following primers: 5’-TGTAAAACGACGGCCAGTTCAAGCATGCAGA GTTGAAG-3’ and 3’-AGGAAACAGCTATGACCATTATGCAAGACACTGGCACTC-5’. The product from PCR2 was sent for sequencing with an M13 universal primer (5’-TTACTTGGCTCAGGTGTGTC-3’). Chromatograms were used to distinguish between WT (TGACTGTTAC**A**GACCCC), *crb2a*^*+/*^- (TGACTGTTAC**A/T**GACCCC) and *crb2a*^-^*/-* (TGACTGTTAC**T**GACCCC).

### Generation of transgenic zebrafish line

The *pTol2 -0.2myl:EGFP-ZO-1* plasmid was generated using Cold Fusion Cloning Kit (System Biosciences, MC101A-SBI). EGFP-ZO-1 was amplified using the Phusion Tag HF Polymerase and the following primers, forward 5’-CAAAGCAGACAGTGAGCTA GCATGGTGAGCAAGGGCGAGGA-3’ and reverse 5’-TCCCCCGGGCTGCAGGAAT TCCTAGAGGCTCGAGATGGGAA-3’. Then the PCR product was cloned into a Tol2 enabled vector with the *-0.2myl7* promoter. After Cold Fusion Cloning colonies containing the construct were sent for sequencing the correct integration of the construct.

### Chemical treatments

Embryos were treated with 10 μM of the ErbB2 inhibitor PD168393 or 10 mM 2,3-butanedione monoxime (BDM) at 54 hpf. In all the treatments, ten embryos per well were placed in 1 mL PTU egg water in 12-well plates containing the corresponding drug. DMSO was used as a control. In the case of PD168393, drugs were added into egg water containing PTU and 1% DMSO to help with solubilization.

### Whole-mount immunostaining of zebrafish embryos/larvae

Embryos were treated with PTU to avoid pigmentation and dechorionated using pronase at 1 dpf. Embryos/larvae were fixed at the stage of interest in 2 ml Fish fix buffer ON at 4°C on nutator. Next day, the Fish fix buffer was removed and embryos were washed 3 × 15 min with PBS/Tween followed by removing of the yolk of the embryos/larvae manually using forceps. Deyolking was followed by permeabilization with Proteinase K (3 μm/ml) in 1 ml PBS/Tween (50-54 hpf → 20 min; 72 hpf → 50 min; 79-80 hpf → 1 hour). After permeabilization embryos/larvae were rinsed quickly with PBDT to stop the proteinase K reaction. Then, embryos/larvae were washed 3 × 15 min in PBDT.

Blocking was done using blocking buffer during 2-3 hours. Then embryos were incubated with the corresponding primary antibody(ies) with Incubation buffer ON at 4°C in the rotator (gently shaking). The following primary antibodies with their corresponding concentrations were used: Crb2a, 1:500 (Zou et al., 2012) was used in Figures 1 and 2; Crb2a, 1:50, (ZIRC, zs-4) was used in the rest of the Figures; ZO1, 1:2000 (Invitrogen 33-9100); EGFP, 1:500 (Aves Lab gfp-1020); α-actinin, 1:1000 (Sigma A7811); N-cadherin, 1:250 (Abcam ab12221); tRFP, 1:500 (Evrogen AB233). Next day, after incubation, the primary antibody(ies) was/were removed and embryos were washed 4 × 20 min with PBDT. Then, embryos were incubated with the secondary antibody(ies) in Incubation buffer for 4 hours at RT in the rotator (gentle shaking). All secondary antibodies were obtained from Life Technologies and used at 1:500. Next, secondary antibody(ies) were removed and embryos were washed first with DAPI in PBS/Tween for 10 min, followed by 6 × 15 min washes in PBS/Tween. All these steps were performed in dark conditions. DAPI (Life Technologies) was used at 1:10.000. Once the washes were finished, embryos were kept in PBS/Tween at 4 °C in darkness until the day of imaging.

### Imaging 36 hpf hearts: embryo preparation

Embryos were treated with PTU to avoid pigmentation and dechorionated using pronase at 1 dpf. At 36 hpf embryos were fixed with 4% PFA ON at 4 °C. Next day, PFA was removed and embryos were washed with PBS/Tween 2 × 10 min. After PFA removal, the heads of the embryos were cut using forceps and a blade. The cut was performed right under the head in an angle of 15 degrees respect the base of the head. Removing the head at early stages help to visualize the heart. Embryos were kept in PBS/Tween at 4°C until the day of imaging.

### Cell transplantation

Donor cells for transplantation were obtained from embryos from *Tg(myl7:MKATE-CAAX); crb2a*^*+/*^- incrosses. WT host embryos were obtained from crossing *Tg(myl7:ras-EGFP)* animals. For transplantation, embryos were first dechorionated by treating with 1 mg/ml pronase in Danieau buffer and embryos were maintained on agarose-coated Petri dishes until 1-cell stage. Embryos were then transferred into 12-well plates where each well is filled with agarose. Being in Ringer’s buffer containing Penicillin/Streptomycin, donor cells were transplanted into 2-3 different WT host embryos along the blastoderm margin at mid-blastula stage. Donor larvae were maintained alive for genotyping. Prior to imaging, selection of mosaic larvae was done by screening for double positive larvae expressing both MKATE-CAAX and ras-EGFP transgenes. Imaging was done using an LSM800 confocal microscope and contribution of the donor-derived cells in the trabecular or compact layer was quantified analyzing plane by plane the whole ventricle using the ZEN software.

### Quantification of trabecular vs compact layer CMs in transplanted hearts

Confocal images were processed and analyzed using ZEN Black Software (Zeiss). The counting was started at the ventricular mid-sagittal plane, counted 12 planes up and 12 planes down at an increment of 1 μm per plane, and used this set of 24 planes for analysis We then counted the number of compact layer and trabecular CMs in the set of 24 planes. Finally, we calculated the percentage of trabecular CMs by dividing the number of trabecular CMs by the total number of CMs.

### Fractional shortening (FS) measurement

Brightfield images of WT, *crb2a*^*+/*^- and *crb2a*^-^*/-* beating hearts were acquired at 48 hpf using Spinning disc confocal microscope. Zebrafish embryos were mounted in low melting agarose containing 0.01% (w/v) tricaine on a glass-bottom dish. Z-stacks of beating hearts were imaged at 400 frames/second. The width of the atrium at the maximum systolic and diastolic phases were measured in three different non-consecutives frames and the FS= (diastole length-sistole length/diastole length) × 100 was calculated.

### In vivo imaging of stopped zebrafish hearts

Embryos and larvae were mounted in 1% low melt agarose on glass-bottom dishes. In order to stop the heart beating before imaging 0.2% (w/v) tricaine was added to the agarose. Z-plane images were taken using a spinning disc confocal microscope (Zeiss, CSU-X1 Yokogawa), LSM800 or LSM880. A 40× (1.1 numerical aperture [NA]) water-immersion objective was used to acquire the images. The optical sections were 1 μm thick.

### Image processing and analysis

Three-dimensional images were processed using Imaris x64 (Bitplane, 7.9.0). To quantify the number of CMs in trabecular versus compact layer CMs, ZEN 2011 software was used. Fiji or Zen softwares were used for the rest of 2D image processing. Images and schemes were prepared using Adobe Photoshop and Power Point.

### Statistical analysis

Data are expressed as mean ± SEM. Differences between groups were compared with a two-tailed Student’s t distribution test. All tests were performed with a confidence level of 95%. (\**P<* 0.05; \*\**P<* 0.01; \*\*\**P<*0.001; \*\*\*\**P<* 0.0001; ns: no significant changes observed).

## Acknowledgments

We would like to thank Xiangyun Wei and Chuanyu Guo for sharing the Crb2a antibody, Elizabeth Knust and Cátia Crespo for providing the *crb2a* mutant and the genotyping protocol, Rubén Marín-Juez, Rashmi Priya, Jason Lai, Felix Gunawan, Sophie Ramas, Chi-Chung Wu and Sally Horne-Badovinac for helpful discussions and sharing reagents and expertise, Rubén Marín-Juez and Rhasmi Priya for critical reading of the manuscript and feedback, Hans-Martin Maischein and Simon Howard for help with zebrafish transplantations and injections, Felix Gunawan for the immunostaining protocol, Rita Retzloff, Martin Laszczyk and their teams for zebrafish care. V.J.A. is a graduate student registered with the Faculty of Biological Sciences at Goethe University, Frankfurt am Main, Germany. Some data from this manuscript form part of V.J.A.’s PhD thesis “Cardiomyocyte apicobasal polarity during cardiac trabeculation in zebrafish”, submitted to Goethe University, Frankfurt am Main, Germany in June, 2018.

## Competing interests

The authors declare no competing or financial interests.

## Author Contributions

V.J.A. and D.Y.R.S. designed the experiments; V.J.A performed all the experiments; V.J.A. and D.Y.R.S. interpreted the data and wrote the paper; D.Y.R.S. supervised the project.

## Funding

These studies were supported by funds from the Max Planck Society.

---

**Figure S1.**
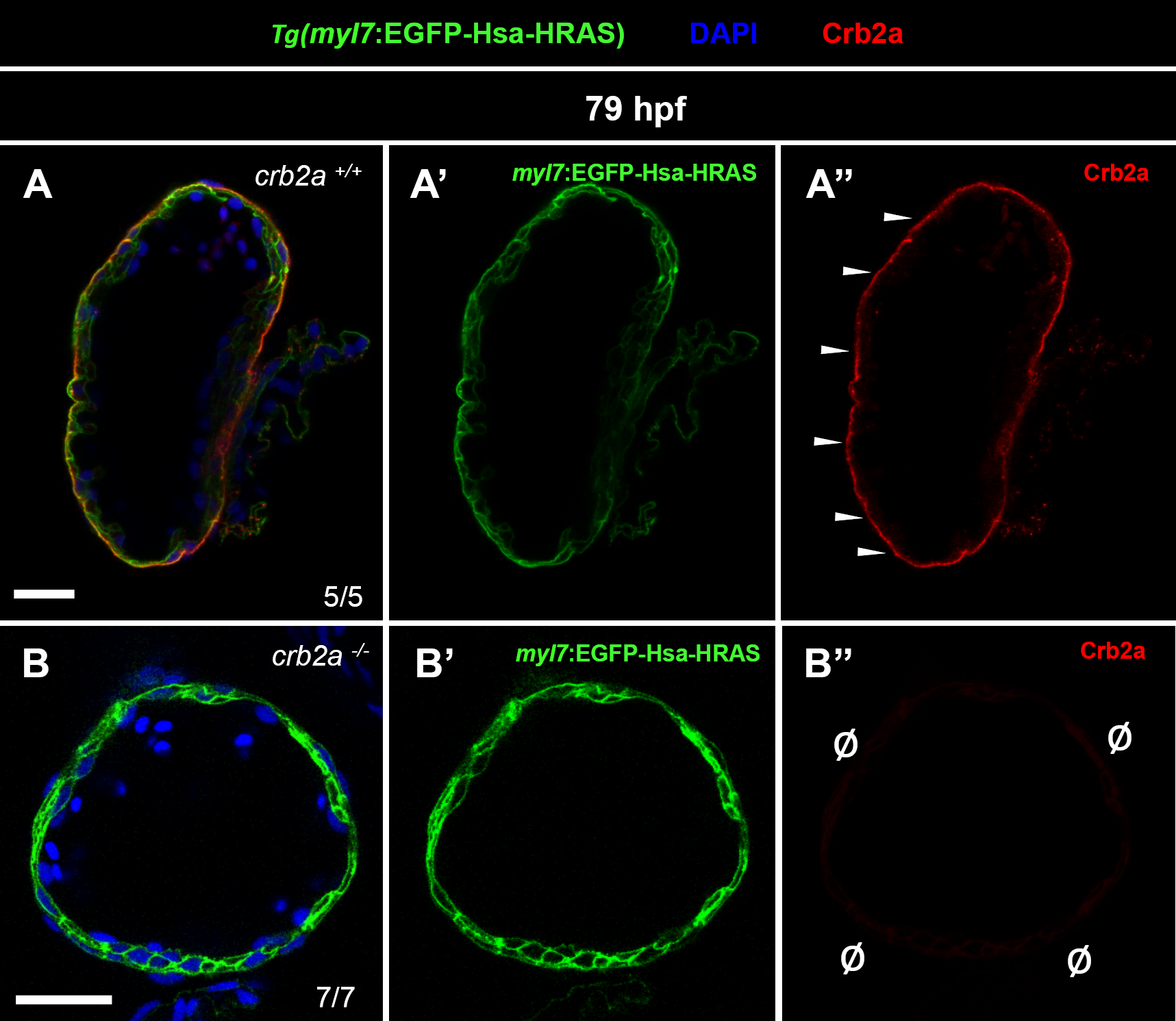
Lack of Crb2a staining in crb2a mutant hearts. (A-B”) Crb2a staining in *79 hpf Tg(myl7:EGFP-Has-HRAS)* hearts of *crb2a*^*+/+*^ (A-A”) and *crb2a*^-^*/-* (B-B”). White arrowheads point to apical localization of Crb2a in *crb2a*^*+/+*^ compact layer CMs (A”), while Ø indicates lack of Crb2a staining in *crb2a*^-^*/-* hearts (B”). V, ventricle. Scale bars, 20 μm.

**Figure S2.**
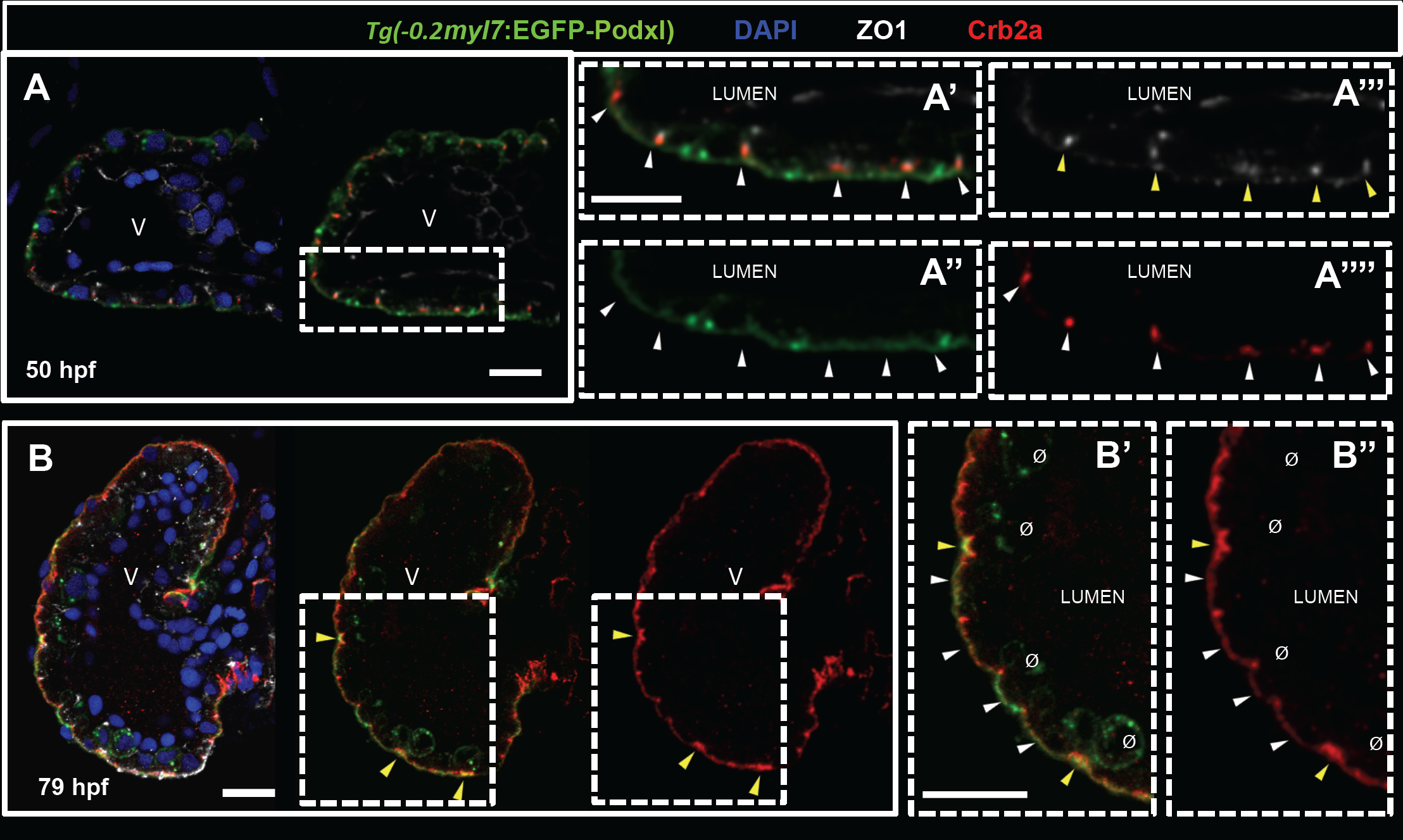
Crb2a co-localizes in early CMs with ZO-1 and subsequently with Podocalyxin. (A-B”) Crb2a immunostaining in *Tg(-0.2myl7:EGFP-podxl)* hearts at 50 (A-A””) and 79 (B-B”) hpf. At 50 hpf, Crb2a localizes to the junctions between CMs at the apical side, coinciding with ZO-1 immunostaining (A’-A””, arrowheads). At 79 hpf, some compact layer CMs undergo apical constriction (B, yellow arrowheads), and Crb2a accumulates at these constricting apical membranes, coinciding with Podocalyxin localization (B’-B”, yellow arrowheads). Moreover, Crb2a expression extends to the apical membrane of compact layer CMs co-localizing with Podocalyxin (B’-B”, white arrowheads). At 79 hpf, Crb2a expression was not observed in delaminated CMs (B’-B”, Ø). V, ventricle. Scale bars, 20 μm.

**Figure S3.**
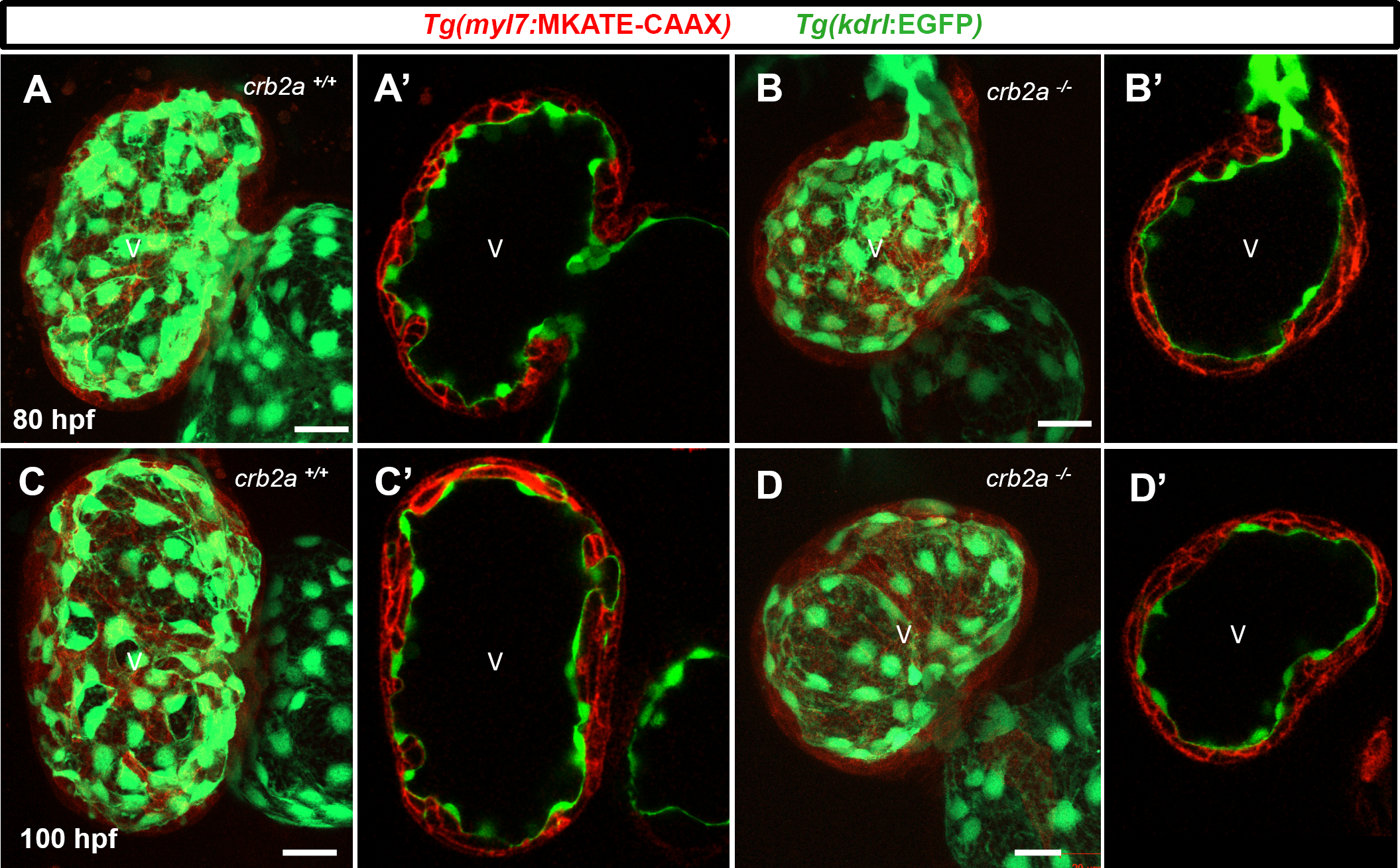
Cardiac jelly reduction is not affected in *crb2a*^-^*/-*. (A-D’) Confocal images (maximum intensity projection) of *Tg(myl7:MKATE-CAAX)*;*Tg(kdrl:EGFP)* hearts at 80 (A-B’) and 100 (C-D’) hpf. *crb2a*^-^*/-* hearts display WT-like reduction of the cardiac jelly at both stages analyzed. V, ventricle. Scale bars, 20 μm.

**Figure S4.**
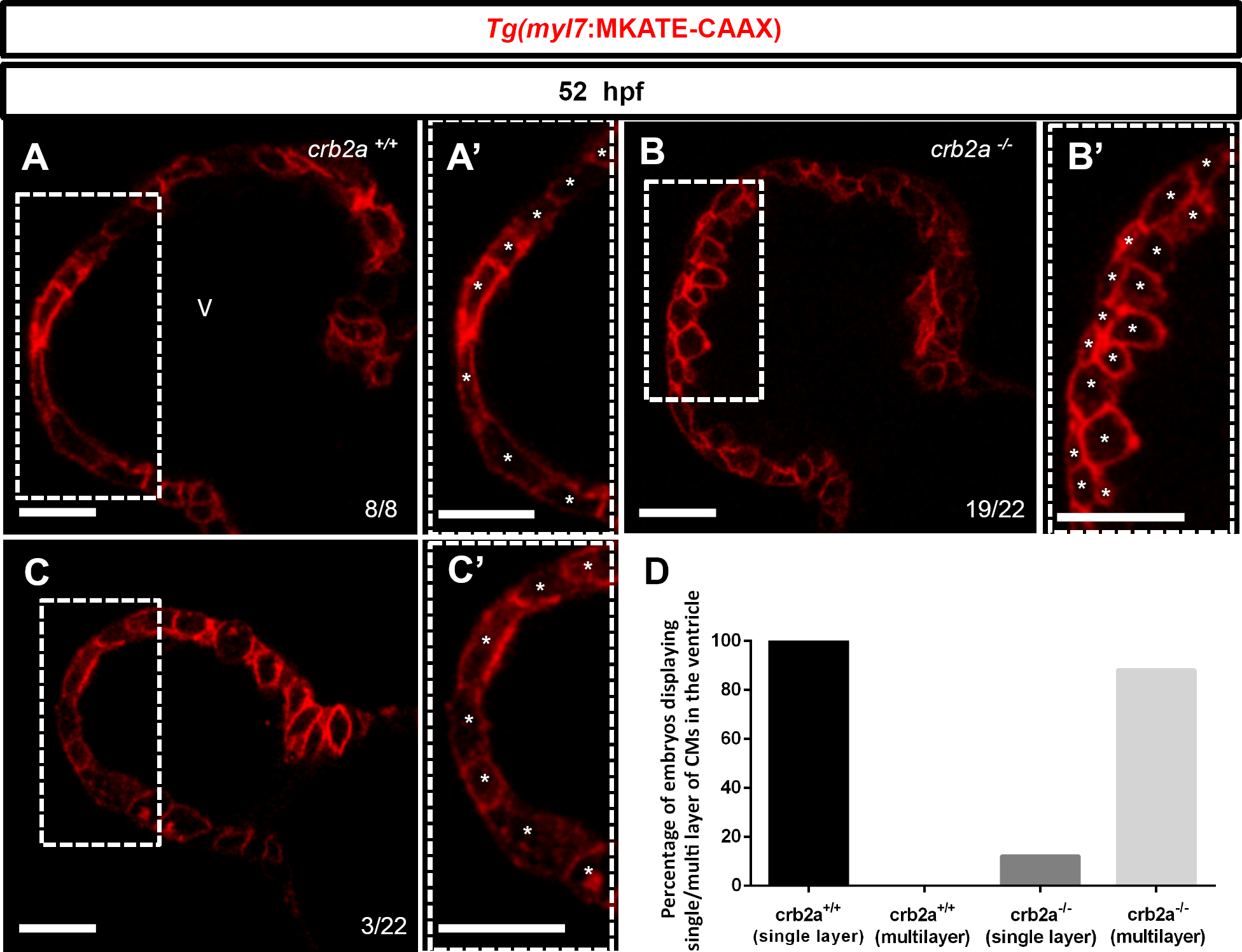
*crb2a*^-^*/-* hearts display CM multilayering at 52 hpf. (A-C’) Confocal images (mid-sagittal sections) of 52 hpf *Tg(myl7:mKATE-CAAX) crb2a*^*+/+*^ (A-A’) and *crb2a*^-^*/-* (B-C’) hearts. Higher magnification images of *crb2a*^*+/+*^ hearts show single layer CMs in the ventricle (A’, asterisks). Higher magnification images show single layer CMs in the ventricle of 3/22 mutants analyzed (B’, asterisks), while the rest of the mutants analyzed display multiple CM layers in their ventricle (C’, asterisks). (D) Percentage of embryos showing single or multiple CM layers in their ventricle. V, ventricle. Scale bars, 20 μm.

**Figure S5.**
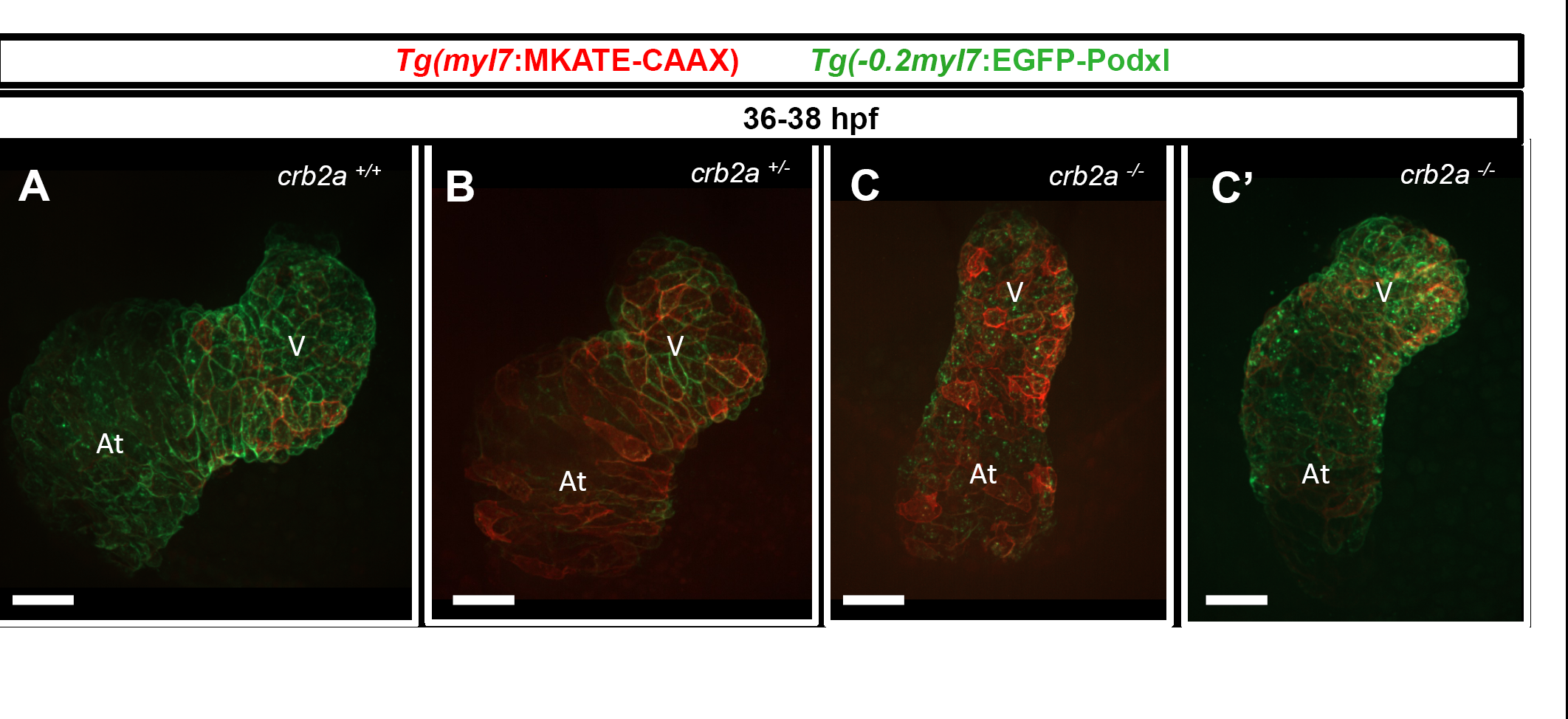
*crb2a* mutant hearts exhibit looping defects. (A-C’) 3D maximum intensity projections of 36-38 hpf *Tg(-0.2myl7:EGFP-podxl)*;*Tg(myl7:MKATE-CAAX*) embryonic hearts. *crb2a*^*+/+*^ and *crb2a*^*+/*^- hearts display complete looping (A (6/6) and B (10/11)), while *crb2a*^-^*/-* hearts display no looping (C (4/11)) or delayed looping (C’ (7/11)). V, ventricle; At, atrium.

**Figure S6.**
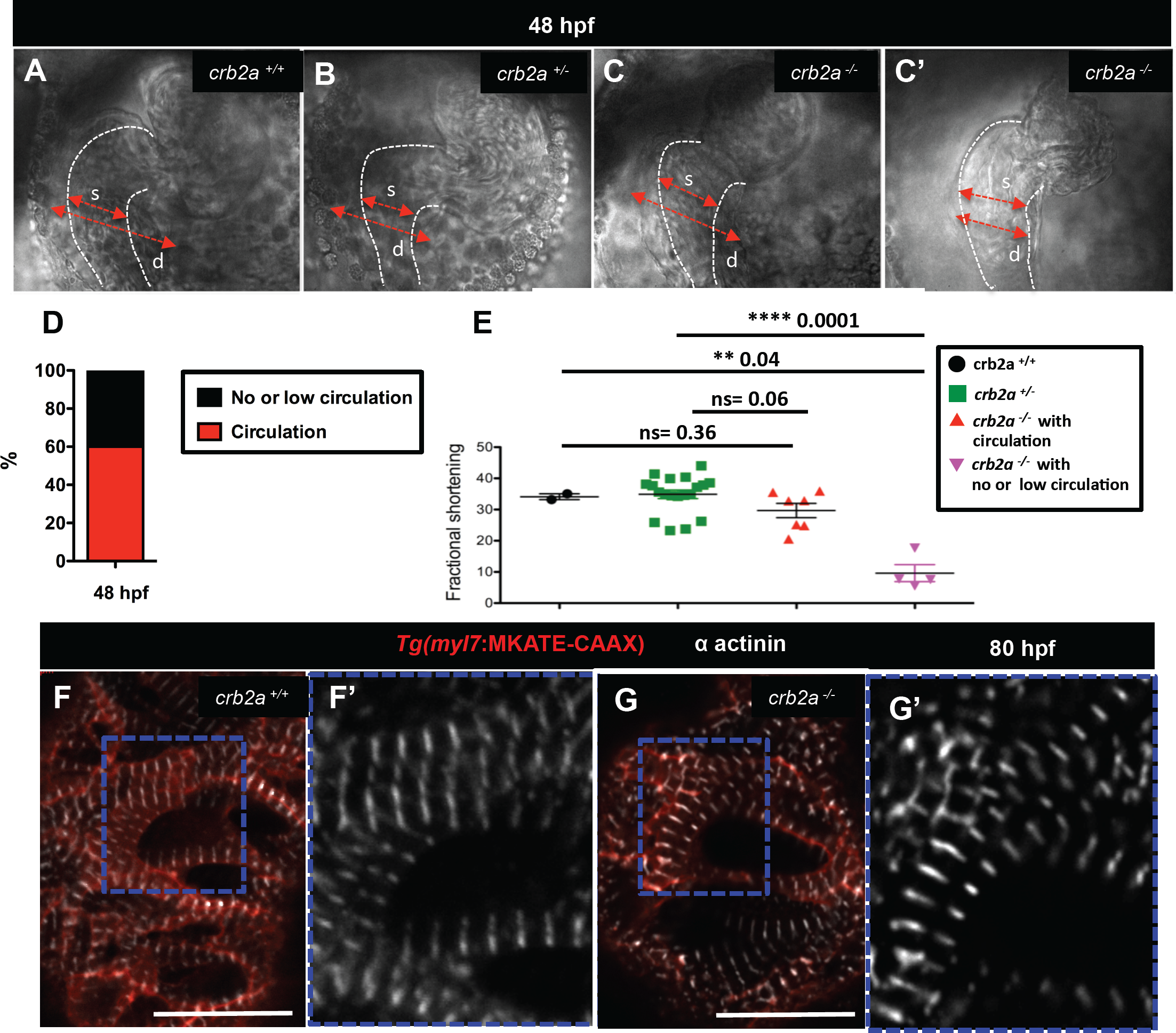
*crb2a*^-^*/-* hearts display slight reduction in contractility at 48 hpf. (A-C’) Brightfield images of 48 hpf embryonic hearts. Red arrows indicate length of diastole (d) and systole (s). White dashed lines represent the heart shape in systole. (D) Percentage of mutant embryos with normal, no, or low circulation at 48 hpf. Total number of embryos analyzed: 650; total number of mutants analyzed, 124 (with circulation (n=50), with low or no circulation (n=74)). (E) Fractional shortening of *crb2a*^*+*^*/+, crb2a*^*+/*^- and *crb2a*^-^*/-*hearts. Data are shown as mean ± SEM. ^**^*P* < 0.001, ^****^*P* < 0.0001 by Student’s t test. (F-G’) Confocal images of α-actinin immunostaining in 80 hpf *Tg(myl7:MKATE-CAAX)* hearts. Blue boxes show high magnification images of the α-actinin immunostaining in *crb2a*^*+/+*^ (F’) and *crb2a*^-^*/-* (G’) hearts. Scale bars, 20 μm.

**Figure S7.**
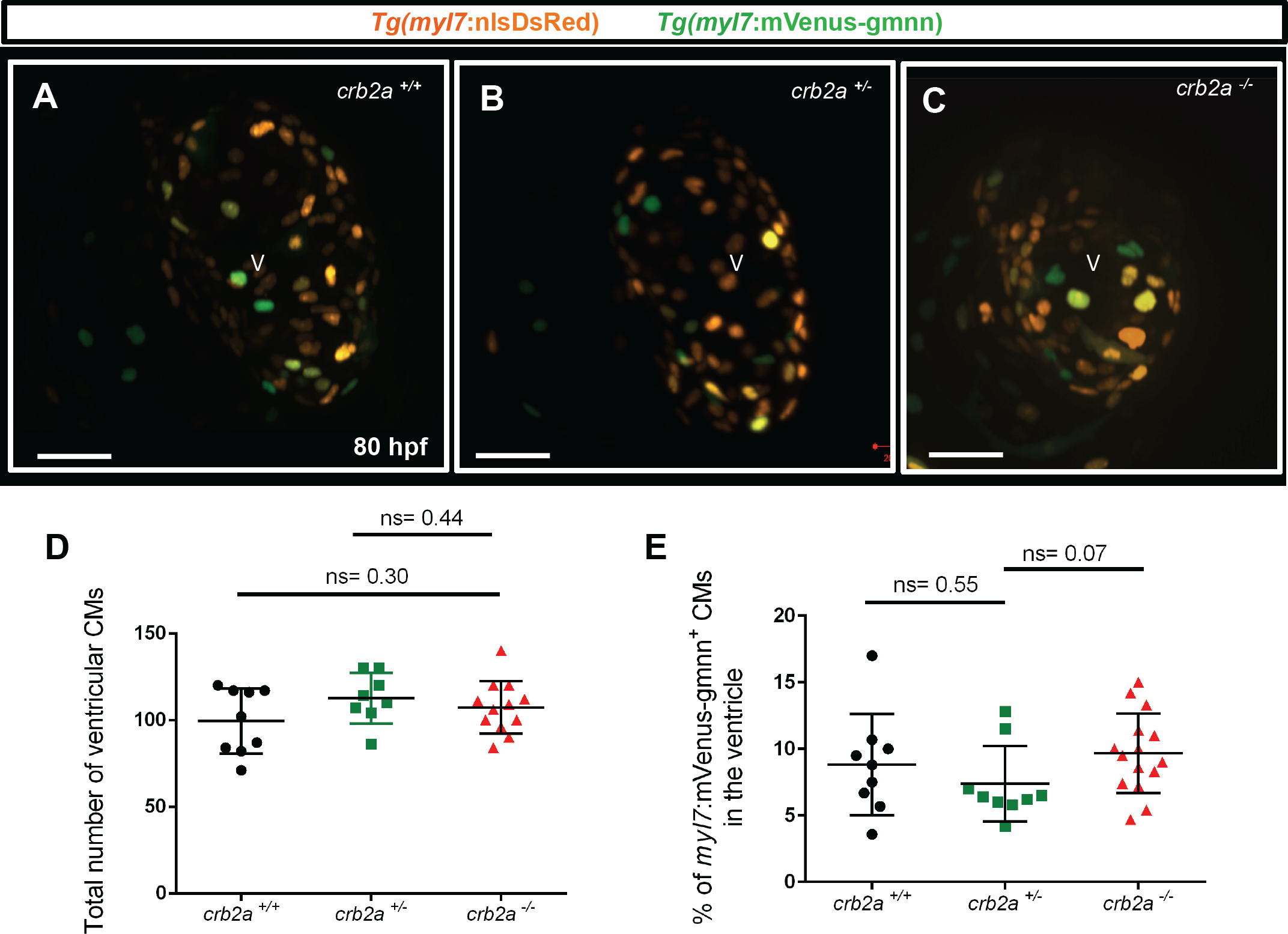
*crb2a*^-^*/-* ventricles do not exhibit an increase in CM proliferation. (A-C) Confocal images (maximum intensity projection) of 80 hpf *Tg(myl7:nlsDsRed); Tg(myl7:VenusGeminin)* hearts. (D) Total number of ventricular CMs. (E) Percentage of *myl7*:mVenus-gmnn^+^ CMs in the ventricle. Each dot represents a heart. Data are shown as mean ± SEM. ns, no significant differences by Student’s t-test. V, ventricle. Scale bars, 20 μm.

**Figure S8.**
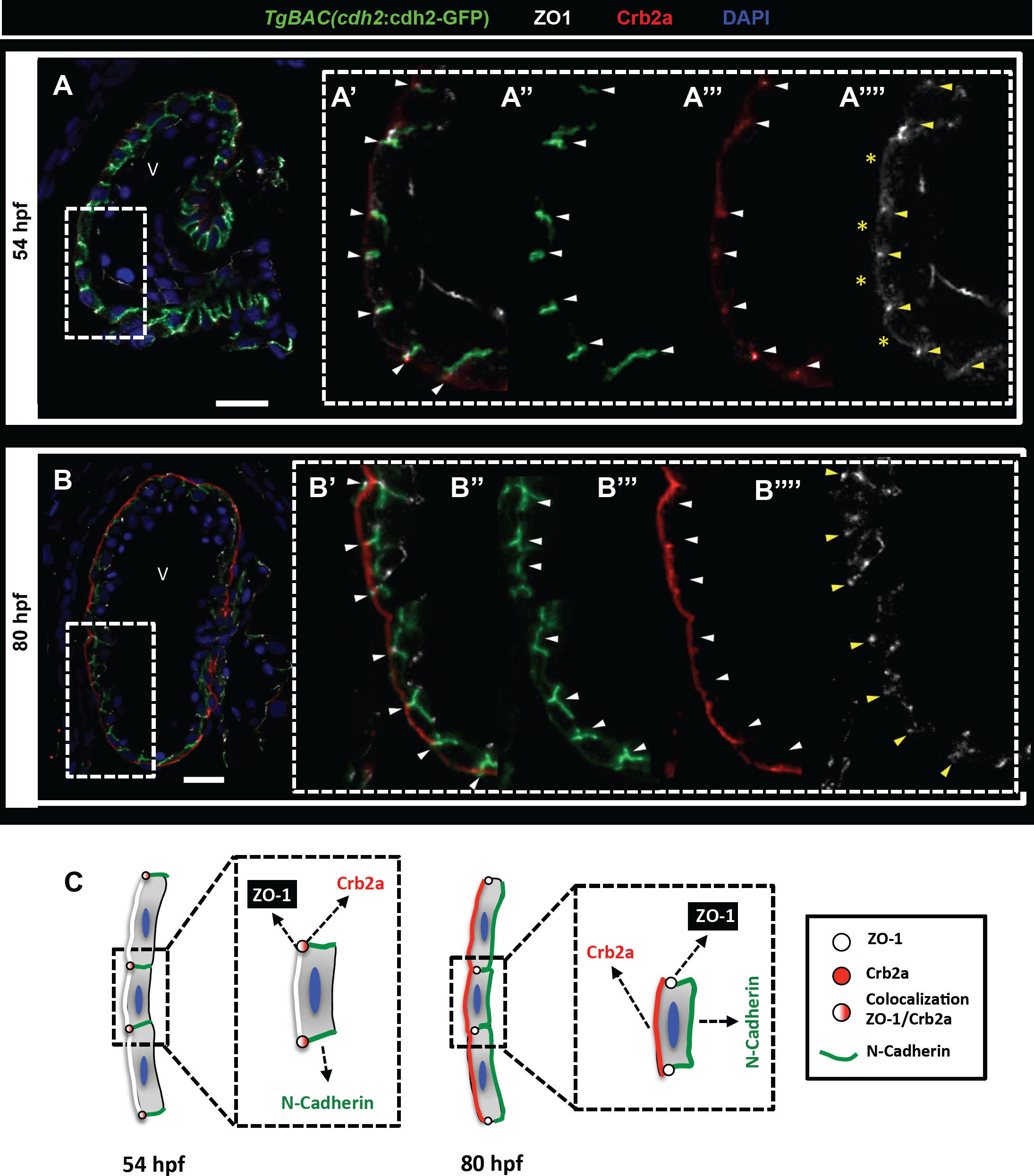
Localization of TJs and AJs, and Crb2a in CMs during trabeculation. (A-B””) Crb2a and ZO-1 immunostainings in *TgBAC(cdh2:cdh2-GFP)* hearts at 54 (A-A””) and 80 (B-B””) hpf. At 54 hpf, white arrows point to lateral localization of N-Cadherin (A’ and A”) and junctional localization of Crb2a (A’ and A”’), yellow arrows and asterisks indicate junctional and apical ZO-1 staining respectively (A””); at 80 hpf, white arrows point to basolateral localization of N-Cadherin (B’ and B”) and apical Crb2a localization (B’ and B”’), yellow arrows point to junctional ZO-1 staining (B””). (C) Schematic representation of tight and adherens junctions in *crb2a*^*+/+*^ CMs. V, ventricle. Scale bars, 20 μm.

**Figure S9.**
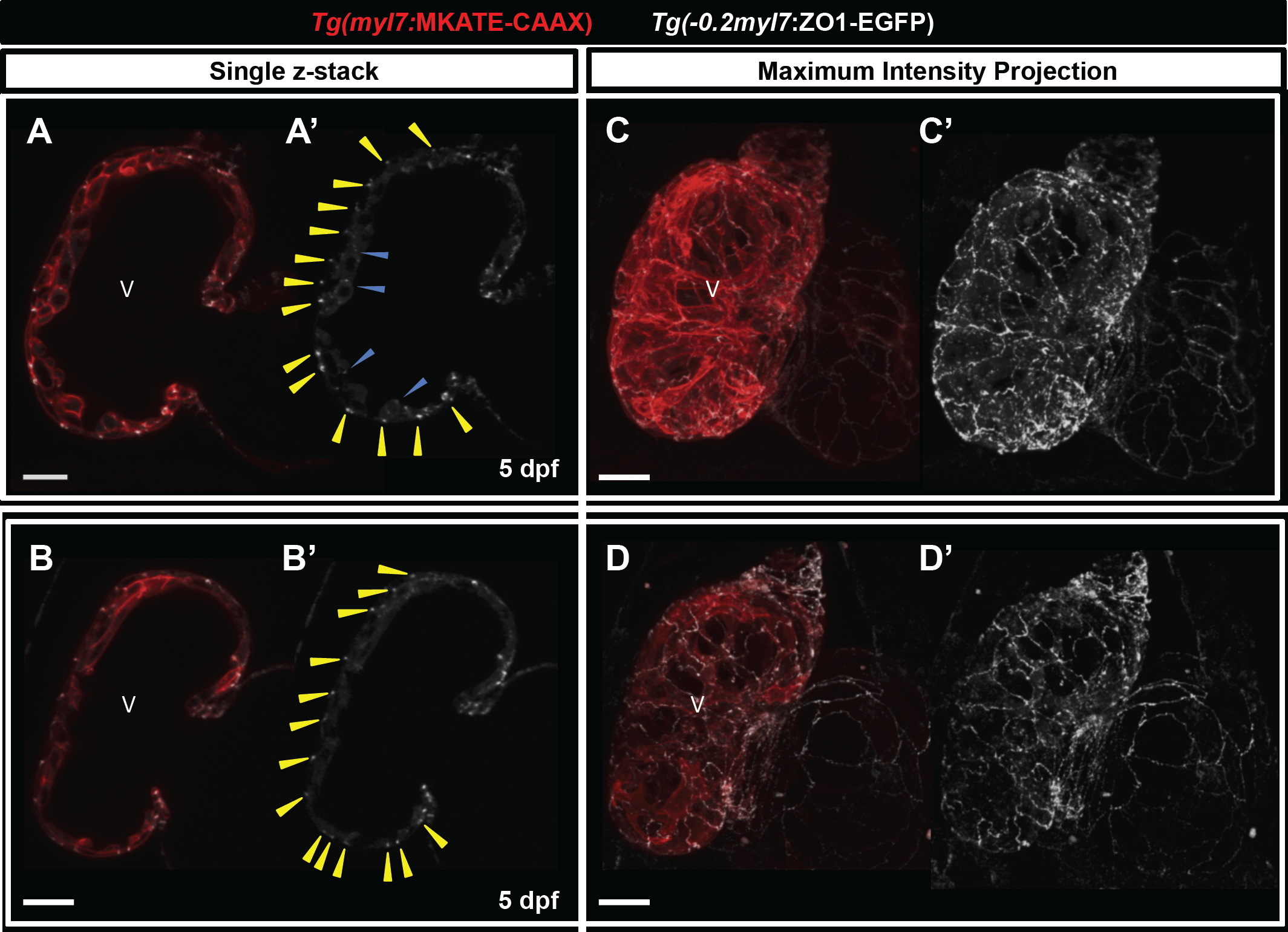
*In vivo* ZO-1 localization in CMs. (A-D’) Confocal images (mid-sagittal sections) (A-B’) and maximum intensity projections (C-D’) of 5 dpf *Tg(myl7:MKATE-CAAX);Tg(-0.2myl7:ZO1-EGFP)* larvae. Yellow arrowheads point to junctional ZO-1 localization in compact layer CMs and blue arrowheads point to junctional ZO-1 localization in trabecular CMs (A’ and B’). V, ventricle. Scale bars, 20 μm.

**Movie S1.** Movie showing Crb2a staining in 3D reconstructed 51 hpf heart

**Movie S2.** Movie showing Crb2a staining in 3D reconstructed 72 hpf heart

**Movie S3**. Movie showing beating of 48-hpf *crb2a*^*+/+*^ zebrafish heart.

**Movie S4.** Movie showing beating of 48-hpf *crb2a*^*+/*^- zebrafish heart.

**Movie S5.** Movie showing beating of 48-hpf *crb2a*^-^*/-* zebrafish heart with normal circulation.

**Movie S6.** Movie showing beating of 48-hpf *crb2a*^-^*/-* zebrafish heart with no or low circulation.

